# Cytoplasmatic polyadenylation of mRNA by TENT5A is critical for enamel mineralization

**DOI:** 10.64898/2025.12.08.692968

**Authors:** Goretti Aranaz-Novaliches, Olga Gewartowska, Frantisek Spoutil, Seweryn Mroczek, Pavel Talacko, Karel Harant, Ana-Matilde Augusto-Vale, Irena Krejzova, Carlos Eduardo Madureira Trufen, Pawel Krawczyk, Ales Benda, Vendula Novosadová, Radislav Sedlacek, Andrzej Dziembowski, Jan Prochazka

## Abstract

Tooth enamel formation, or amelogenesis, is a complex biomineralization process regulated by ameloblasts cells. These cells secrete enamel matrix proteins (EMPs), including Amelogenin (AMELX) and Ameloblastin (AMBN), which are essential for hydroxyapatite crystal deposition during enamel mineralization. Precise regulation of this process ensures the mechanical and chemical strength of dental enamel. Using Tent5a knock-out (KO) mouse, we demonstrated that TENT5A, a non-canonical poly(A) polymerase is crucial for the stability, translation, and secretion of EMP mRNAs, particularly AMELX, during amelogenesis. TENT5A-deficient mice exhibited Amelogenesis imperfecta, characterized by enamel hypomineralization, reduced thickness, and disrupted ultrastructure, as revealed by micro-computed tomography. Through nanopore direct mRNA sequencing, we identified that TENT5A polyadenylates *Amelx* and other mRNAs encoding secreted proteins, enhancing their stability and translation in the endoplasmic reticulum. Moreover, loss of TENT5A altered AMELX secretion and extracellular self-assembly, impairing matrix organization and hydroxyapatite deposition. This study highlights the role of TENT5A in post-transcriptional regulation during enamel formation, demonstrating its importance in enamel homeostasis.

**Teaser:** TENT5A adds poly(A) tails to specific mRNAs in enamel, enhancing their expression, which is crucial for dental mineralization.

## Introduction

Eukaryotic gene expression is regulated at multiple levels, and post-transcriptional mRNA modifications play a crucial role. In addition to canonical mRNA polyadenylation, which occurs in the nucleus, there are non-canonical poly(A) polymerases (ncPAPs) that function in the cytoplasm by adding an extra poly(A) tail to selected mRNAs. These ncPAPs belong to the terminal nucleotidyl transferases (TENTs) superfamily and their role is to protect mRNA from degradation, confer stability, and promote mRNA translation (*1–4*).

TENTs have been shown to play a role in embryonic germline and embryonic development by promoting translation of dormant maternal mRNA during zygote genome activation (*5*, *6*). Furthermore, are critical for somatic cell homeostasis (*7*, *8*), for example, TENT2 promotes synaptic plasticity in neurons while TENT5C is involved in antibody production in T cells and suppresses multiple myeloma progression (*2*, *9*, *10*).

Among the TENT proteins is TENT5A, formerly known as FAM46A, which belongs to the TENT5 protein family (TENT5A, TENT5B, TENT5C and TENT5D). TENT5A has been associated with various human diseases such as retinitis pigmentosa and osteogenesis imperfecta (OI) (*11*, *12*). In our previous studies, we generated a Tent5a knock-out (KO) mouse model displaying OI disease. We further demonstrated that TENT5A is a cytoplasmic ncPAP that selectively adds an additional poly(A) tail to mRNAs, thereby stabilizing transcripts (*13*, *14*). Furthermore, we found that TENT5A is expressed in osteoblasts and directly polyadenylates collagen and other extracellular protein transcripts, enhancing their translation during bone formation.

In this study, we investigate the role of TENT5A in tooth enamel formation. While both bones and teeth are mineralized tissues, the processes governing their development are notably distinct. Unlike bones, enamel is a highly structured tissue composed almost entirely of inorganic hydroxyapatite. Due to its developmental origin and biochemical properties, enamel lacks the capacity for self-renewal or repair. Therefore, the molecular driving enamel development and maturation must be tightly regulated to ensure the creation of a structure capable of maintaining lifelong tooth function.

We discovered that Tent5a KO mice exhibit hypomineralization of the teeth and reduction of the enamel layer, revealing the role of ncPAPs in enamel formation for the first time. Amelogenesis, the process of enamel formation, is coordinated by ameloblast epithelial cells, which are crucial during the secretory stage as they synthesize and secrete enamel matrix proteins (EMPs), primarily amelogenin (AMELX) and ameloblastin (AMBN). This process results in the formation of an extracellular organic matrix composed of 90% AMELX and 10% AMBN, along with other minor EMPs (Sire et al., 2007a). This organic matrix provides a scaffold that promotes the oriented crystallization of hydroxyapatite crystals (HAPs), ultimately leading to enamel mineralization (*16*).

Although the production and extracellular interaction between AMELX and AMBN is critical for proper enamel formation, the precise regulation of this process remains elusive (*17–19*). Comprehensive analysis of Tent5a KO mice revealed a disruption in enamel microstructure and altered self-assembly of AMELX and AMBN in the extracellular organic matrix. Furthermore, TENT5A was localized in the endoplasmic reticulum (ER) of ameloblast cells where it promotes the expression of *AmelX* and other secreted protein transcripts. Notably, AMELX, a substrate of TENT5A, and AMBN, which is not, exhibited altered secretory patterns in Tent5a KO ameloblasts. In conclusion, this study provides the first evidence that modulation of poly(A) tail length by TENT5A is crucial for coordinating the synthesis and proper secretion of AMELX and other mineralization-supporting genes in ameloblasts. This regulation is essential for proper organic matrix formation, HAP deposition, and enamel mineralization.

## Results

### *Tent5a* ablation causes amelogenesis imperfecta

In our previous study, we examined the phenotype of a Tent5a KO mouse model (*14*), which exhibited hypomineralized bones. Despite differences in the origin and biomineralization mechanism between bone and enamel, we observed significant structural and mineralization defects in the teeth of Tent5a KO mice. X-ray absorption intensities revealed that Tent5a KO mice had hypomineralized teeth, often resulting in premature abrasion or fracture. Additionally, generalized hypomineralization of the cranial bone was observed. Micro-computed tomography (μCT) phase-contrast visualization showed a reduction in the enamel layer, as indicated by the pseudocolor scheme, suggesting a disruption in the amelogenesis process (Fig 1a).

**Fig. 1.**
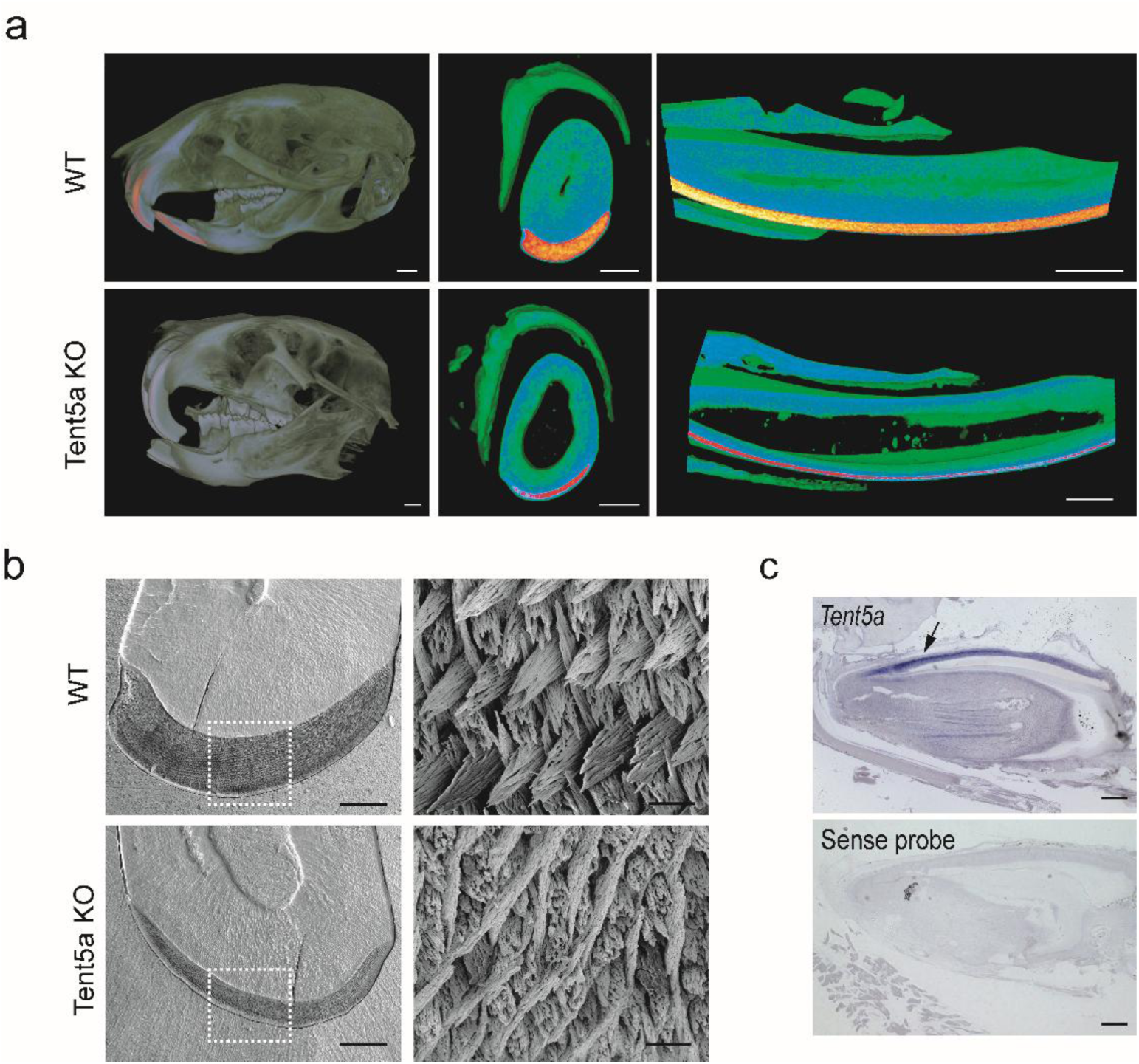
Tent5a KO mice displays amelogenesis imperfecta and Tent5a expression in teeth. (**A**) 3D-micro-computed tomography (µCT) images of representative WT (up row) and Tent5a KO mice (lower row). Mouse head showing broken and hypomineralized incisor and phase contrast μCT images of transversal and Longitudinal sections of mouse incisor teeth. Green-to-yellow pseudocolor bars represent the increasing level of mineralization. Scale bar, respectively, 1mm, 250μm and 500μm. (**B**) SEM images of a mouse incisor cross-section displaying WT and Tent5a KO enamel. Enamel thickness and crystallite organization is altered in Tent5a KO samples. Scale bar, respectively, 100μm and 2.5μm. (**C**) In situ hybridization of *Tent5a* mRNA showing expression in secretory ameloblasts in mouse incisor teeth. Scale bar, 200µm.

Scanning electron microscopy (SEM) revealed a thin enamel layer with a disorganized crystal structure and a weak hydroxyapatite (HAP) crystallite composition in the enamel of Tent5a KO mice. Moreover, there was an increase in the amount of interprismatic matrix (IPM) in Tent5a KO enamel, leading to a reduction in prism volume (Fig 1b). Taken together, these results suggest that TENT5A plays an essential role in regulating HAP deposition and further enamel maturation. While enamel is not completely absent in Tent5a KO mice, its structural integrity is severely compromised, resulting in a condition known as amelogenesis imperfecta (AI) type IV. This type of AI causes hypomaturation-hypoplastic enamel, characterized by thin enamel with areas that are hypomineralized (*20*, *21*).

Despite enamel hypomineralization, the morphology of ameloblast cells in Tent5a KO incisors was normal, and the location and cell polarity of secretory stage ameloblast were indistinguishable from those of the WT teeth (Fig S1). Furthermore, *Tent5a* mRNA was specifically expressed in secretory ameloblasts, which are responsible for secreting enamel matrix proteins (EMPs) essential for enamel formation (Fig 1c). These findings are consistent with a single-cell transcriptomics study from mouse incisors (*22*) which identified *Tent5a* expression in distal secretory ameloblasts. Therefore, deletion of Tent5a has an autonomous effect on ameloblasts and potentially secretory function during enamel formation.

### *AmelX* mRNA polyadenylation by TENT5A is critical for RNA stability and protein synthesis

Since the organic matrix of enamel is mainly formed by amelogenin and ameloblastin, we investigated their gene expression in ameloblasts of Tent5a KO mice. Accordingly, qPCR analysis of incisor ameloblasts (proximal part of the differentiating ameloblast) revealed a 2-fold decrease (p-value < 0.01) in *AmelX* expression compared to WT (Fig 2a). This decrease in mRNA expression was accompanied by a reduction in amelogenin protein synthesis in teeth (Fig 2b). However, the expression of ameloblastin, another relevant EMP protein, showed a trend towards lower expression in Tent5a KO mice, although it did not reach statistical significance (Fig 2a, 2b & Fig S2a).

**Fig. 2.**
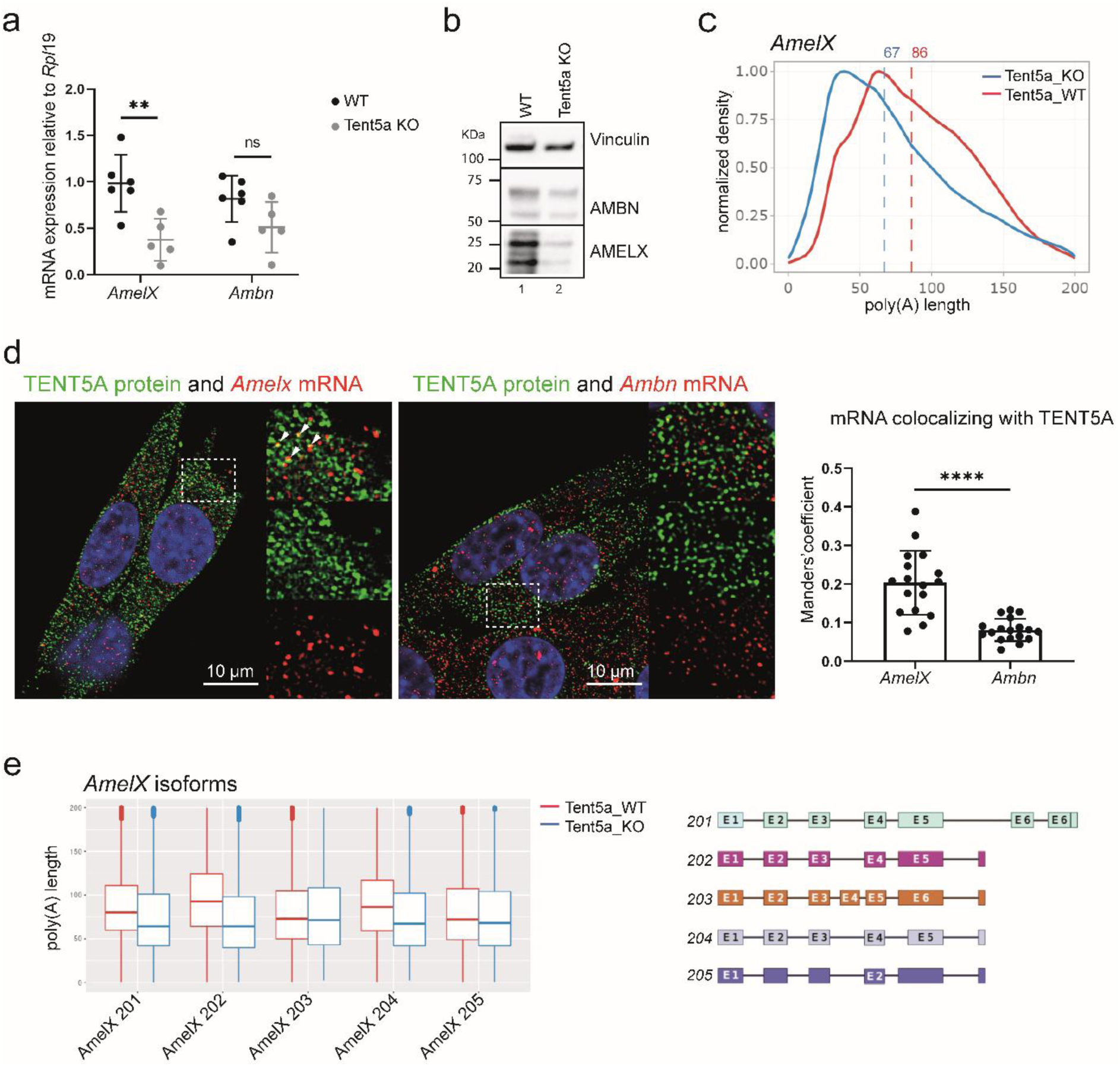
*AmelX* mRNA is polyadenilated by TENT5A. (**A**) mRNA levels of *AmelX* and *Ambn* in mice ameloblasts normalized to *Rpl19* measured by RT-qPCR. Data bars are mean±SEM. Statistical tests are unpaired student t-tests, *p-value <0.05, **p-value <0.01. (**B**) Western blot analysis of AMELX and AMBN protein levels in WT (lane 1) and Tent5a KO (lane 2) teeth. (**C**) Direct RNA sequencing (DRS)-based poly(A) length profiling of *AmelX* mRNA isolated from incisor cervical loop, p-value < 0.001. (**D**) TENT5A protein and *AmelX* and *Ambn* mRNA localization in LS8 ameloblast-like cells. Scale bar, 10 μm. Colocalization parameters were measured using Manders’ coefficient, statistical test Non-parametric Paired Wilcoxon, ****p-value < 0.0001. (**E**) All *AmelX* isoforms scheme and comparison of poly(A) tail length in WT and Tent5a KO ameloblast measured by Nanopore DRS.

To investigate whether TENT5A directly targets *AmelX* mRNA for polyadenylation, we conducted direct mRNA sequencing (DRS) and found that *AmelX* transcripts in Tent5a KO samples had a shorter poly(A) tail compared to WT samples (WT=86 vs KO=67, p>0.001) (Fig 2C). Moreover, *Ambn* transcript showed a slightly longer poly(A) tail in Tent5a KO ameloblasts (WT=80 vs KO=88, p>0.001) (Fig S2b). Thus, *AmelX* transcript, but not *Ambn*, is a substrate of TENT5A. Additionally, the colocalization of TENT5A with *AmelX* mRNA but not *Ambn* was confirmed by the mRNA FISH experiment. The analysis revealed a significant difference in colocalization, with *AmelX* exhibiting a Manders’ coefficient of 0.2 compared to 0.08 for *Ambn* (Fig 2d).

Further analysis revealed that TENT5A preferentially targets specific *AmelX* transcript variants. Among them, ENSMUST00000066112.12 (Amelx-201), ENSMUST00000112118.8 (Amelx-202) and ENSMUST00000112120.8 (Amelx-204) showed the most significant decrease in poly(A) tail length. All *AmelX* transcript variants were transcribed in ameloblasts, with their expression reduced in Tent5a KO samples. Notably, Amelx-202 showed the highest expression in ameloblasts and the largest difference in poly(A) tail length (−29A). In contrast, *AmelX* transcripts ENSMUST00000112119.8 (Amelx-203) and ENSMUST00000154923.2 (Amelx-205) showed no significant differences in poly(A) tail length and had the lowest expression levels. Notably, Amelx-203 includes an additional exon 4, absent in other *AmelX* transcripts, while Amelx-205 is not annotated as a protein-coding gene (Fig 2e & S2c).

To study whether TENT5A targets specific *AmelX* splice variants based on their sequences, we conducted an RNA stability analysis in the immortalized mouse ameloblast-like (IMAL) cells, which expresses *Tent5a* but lacks endogenous *AmelX*. We transfected in-vitro transcribed Amelx-202 (a TENT5A substrate) and Amelx-203 (a non-substrate) mRNA into the cells, hypothesizing that Amelx-202 would show greater persistence. Indeed, Amelx-202 mRNA demonstrated higher stability compared to Amelx-203 mRNA. Interestingly, comparing in-vitro synthesized Amelx-202 mRNA with and without a poly(A) tail revealed no difference in stability. However, in the case of Amelx-203, artificially adding a poly(A) tail before transfection had a stabilizing effect (Fig S2d). This suggests that TENT5A activity is not dependent on the canonical poly(A) tail and may instead involve a more context-specific mechanism.

### AMELX secretion and extracellular assembly are altered in Tent5a KO mice

Next, we examined the effect of Tent5a deficiency on the secretion and self-assembly of AMELX and AMBN within ameloblasts. In the extracellular organic matrix, AMBN and AMELX self-assemble and interact to form the enamel matrix, which is essential for directing hydroxyapatite deposition and subsequent mineralization (*23*). In the absence of TENT5A, the thickness and integrity of the enamel matrix was reduced. Although AMELX and AMBN were present in the extracellular matrix, they were unable to assemble into a precisely organized mesh-like structure as observed in WT controls. (Fig 3a).

**Fig. 3.**
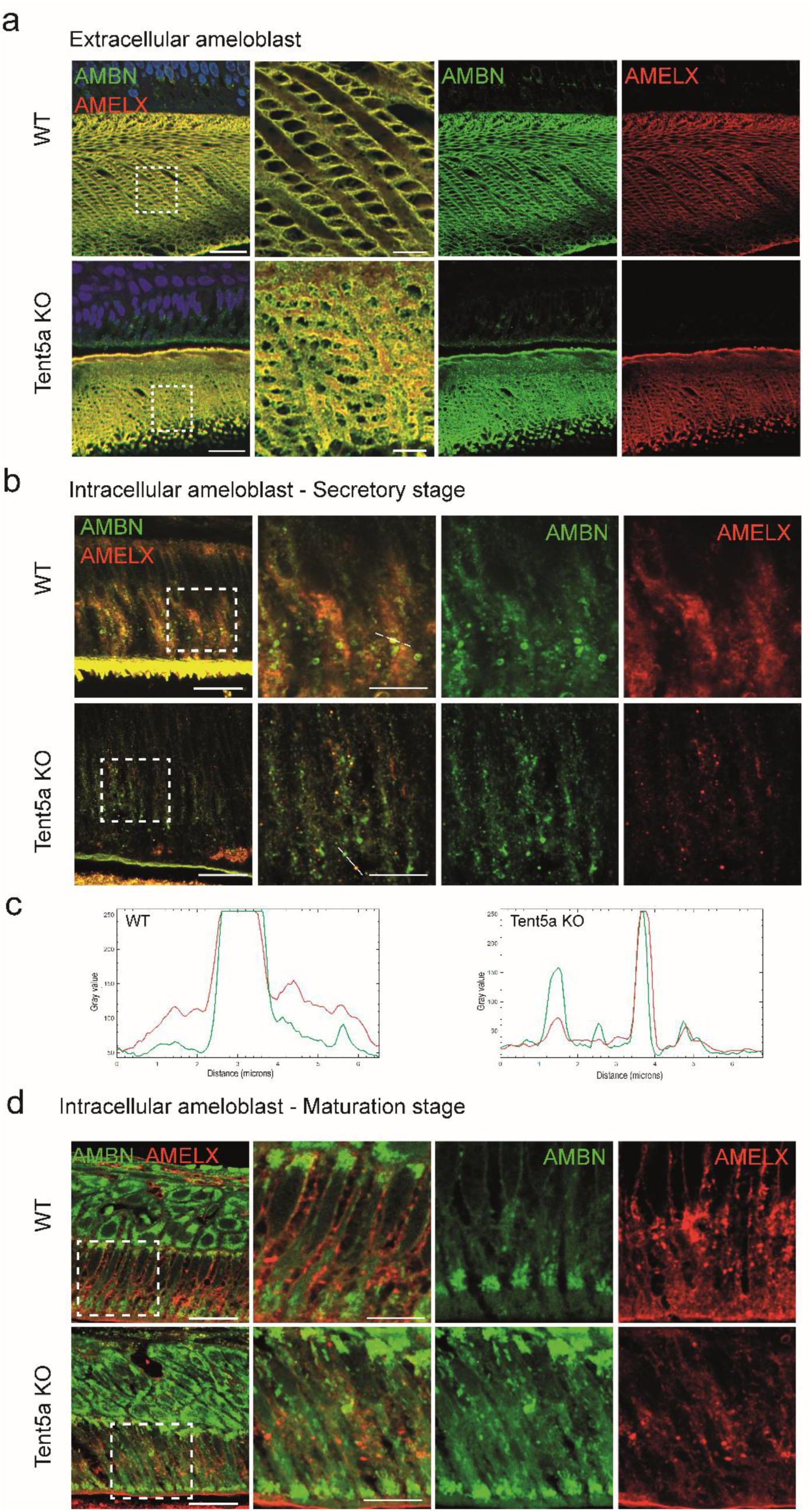
Immunostaining of AMELX (red) and AMBN (green) in WT and Tent5a KO mouse ameloblasts. (**A**) Enamel organic matrix in the extracellular region of maturation stage ameloblasts and high magnification image of enamel. Scale bar, respectively, 20μm and 5μm. (**B**) Intracellular region of secretory stage ameloblasts cells. Scale bar, respectively, 20μm and 10μm. (**C**) Histogram of colocalization of AMELX and AMBN signal intensities in secretory vesicles. Quantification was measured from the straight line marked in intracellular ameloblasts. (**D**) Maturation stage intracellular region of ameloblasts. Scale bar, respectively, 20μm and 10μm.

Furthermore, we observed changes in the secretion dynamics of AMELX and AMBN. In WT ameloblasts during the secretory stage, AMELX was consistently localized in broadly distributed intracellular secretory vesicles as well as the extracellular matrix. Conversely, AMBN showed an exclusive presence within well-defined vesicles, occasionally accompanied by a more diffuse pattern alongside AMELX. In contrast, the deficiency of TENT5A led to a reduction of distinct AMBN vesicles, causing both proteins to primarily colocalize within smaller secretory vesicles. This shift was particularly distinct for AMELX, which showed a significant reduction in production, as detected by immunofluorescence, with a lesser decrease observed for AMBN (Fig 3b, c, d).

During the maturation stage of ameloblasts, AMBN was observed in both the basal and secretory poles of the cells. This distribution remained consistent in Tent5a KO ameloblasts, although the intracellular distribution was more diffuse. In contrast, AMELX, showed a uniform localization in the apical region and near the nucleus in WT mice, showed a significant reduction in Tent5a KO mice, and was exclusively localized in the mid-apical region of ameloblasts (Fig 3e). These findings suggest that Tent5a deficiency leads to alterations in the secretory pathway and extracellular self-assembly of enamel matrix proteins. Ultimately, this affects the processes of enamel deposition and mineralization.

### TENT5A prefers to polyadenylate mRNAs encoding secreted proteins which are involved in mineralization

We asked whether the role of TENT5A in ameloblasts is specific for post-transcriptional regulation of AmelX or has a general function of mRNA regulation. A bulk transcriptomic assay revealed that Tent5a KO ameloblasts showed significant downregulation of several secreted proteins important for proper amelogenesis. Although a similar number of upregulated genes (48%) and downregulated genes (52%) were identified, the downregulated genes had a lower p-value when ordered by statistical significance (Fig 4a). The top downregulated genes included *AmelX*, collagen (*Col1a1*), biglycan (*Bgn*) and clusterin (*Clu*), with corresponding log2 fold changes, −2.45, −1.78, −1.12, −1.68 (Fig S3a).

**Fig. 4.**
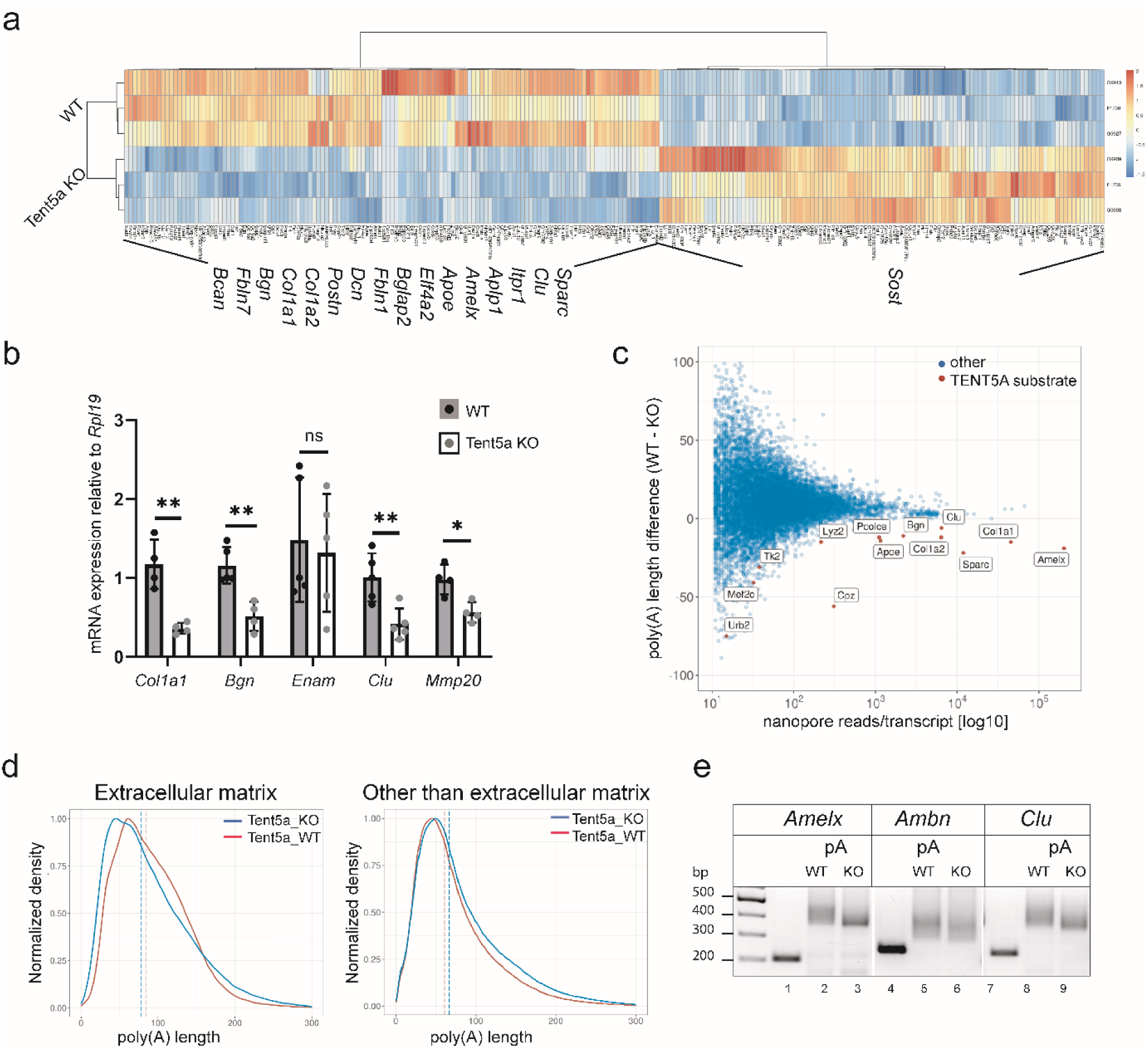
Identification of TENT5A substrates in ameloblasts by transcriptomics assays. (**A**) Illumina sequencing mRNA transcriptomics heatmap comparing WT and Tent5a KO cervical loop samples. Relevant dysregulated genes for Amelogenesis are presented. N=3, p-value <0.05. (**B**) RT-qPCR analysis of enamel proteins in mice ameloblasts normalized to *Rpl19*. Data presented as mean ± SD. Statistical tests are unpaired student t-tests, *p-value p<0.05, **p-value p<0.01. (**C**) DRS of WT and Tent5a KO cervical loop RNA. MA-like plot showing transcripts with a significant difference in poly(A) tail length. Adjusted p-value < 0.05, length difference >5, at least 10 reads in WT sample. (**D**) Distribution of poly(A) tail length of mRNAs encoding extracellular matrix and other proteins in WT and Tent5a KO ameloblasts. (**E**) A PCR-based PAT assay shows the poly(A) tail length of *AmelX*, *Ambn* and *Clu* transcripts in ameloblasts. Lanes 1, 4 and 7 are gene-specific PCR products, lanes 2, 5 and 8, WT ameloblast poly(A) tail PCR product and lanes 3, 6 and 9 Tent5a KO samples.

Finding similar expression patterns to those observed in the RNAseq datasets, qPCR analysis confirmed significantly lower mRNA expression of *Clu, Col1a1, Bgn*, and *Mmp20* in Tent5a KO tissue. In contrast, the enamel matrix protein enamelin (*Enam*) showed no significant difference in expression (Fig 4b), suggesting that, similar to *Ambn, Enam* mRNA is not regulated by TENT5A. Clusterin protein levels corresponded to mRNA quantification and confirmed the decreased protein production of the affected transcript (Fig S3b).

Then, the abovementioned DRS analysis was expanded to perform a genome-wide poly(A) profiling assay to identify all potential TENT5A substrates in ameloblasts. First, the global poly(A) tail distribution was checked, and it was confirmed that it was not affected; therefore, TENT5A does not affect all mRNAs, only selected transcripts (Fig S3c). We then examined the differences in poly(A) tails between WT and Tent5a KO ameloblast, which revealed 14 mRNAs with significantly shorter tails in the Tent5a KO group (Fig S3d). These mRNAs were also the most downregulated genes and thus are likely to be stabilized by TENT5A-mediated polytadenylation. Alongside the previously identified *AmelX* transcript, *Col1a1, Clu, Col1a2*, and *Bgn* were identified as TENT5A substrates (Fig 4c). Specifically, the median poly(A) tail length from *Clu* was reduced from 82nt in WT to 76nt in KO, whereas *Bgn* was decreased 91nt in WT to 80nt in KO (Fig S3d). All these genes are extracellular matrix proteins that are post-transcriptionally regulated by TENT5A. Notably, *Mmp20* and *Enam* were not identified as TENT5A substrates. When further dividing the DRS analysis into extracellular matrix (ECM) transcripts and non-ECM transcripts, there was a difference in the poly(A) tail length (Fig 4d).

The individual putative substrates of TENT5A were individually validated by poly(A) profiling (PAT) assay (Fig 4e). Together, these experiments indicated that *AmelX* is the major target of TENT5A but the downregulation of other extracellular proteins, such as extracellular protein chaperone clusterin and the structural proteoglycan biglycan, may have an additive effect on the hypomineralized enamel phenotype. The complexity of amelogenesis requires the coordination of multiple extracellular matrix proteins and TENT5A is a key regulator of their expression.

### TENT5A functions in the ER membrane and promotes enamel extracellular proteins synthesis

To investigate the subcellular localization of TENT5A and its role in ECM regulation during amelogenesis, we analysed its localization in mouse ameloblast cells. We found that TENT5A was more enriched in the cytoplasmic fraction compared to the nuclear fraction (Fig S4). We confirmed this cytoplasmic localization through immunofluorescence assays conducted on IMAL and LS8 ameloblast-like cells (*24*). Additionally, some areas showed colocalization of TENT5A with calnexin ER protein (Fig 5a). Nevertheless, subcellular localization alone does not determine substrate specificity. While TENT5A does not regulate all secreted proteins, such as ameloblastin, it appears to play a significant role in regulating certain proteins critical for enamel formation.

**Fig. 5.**
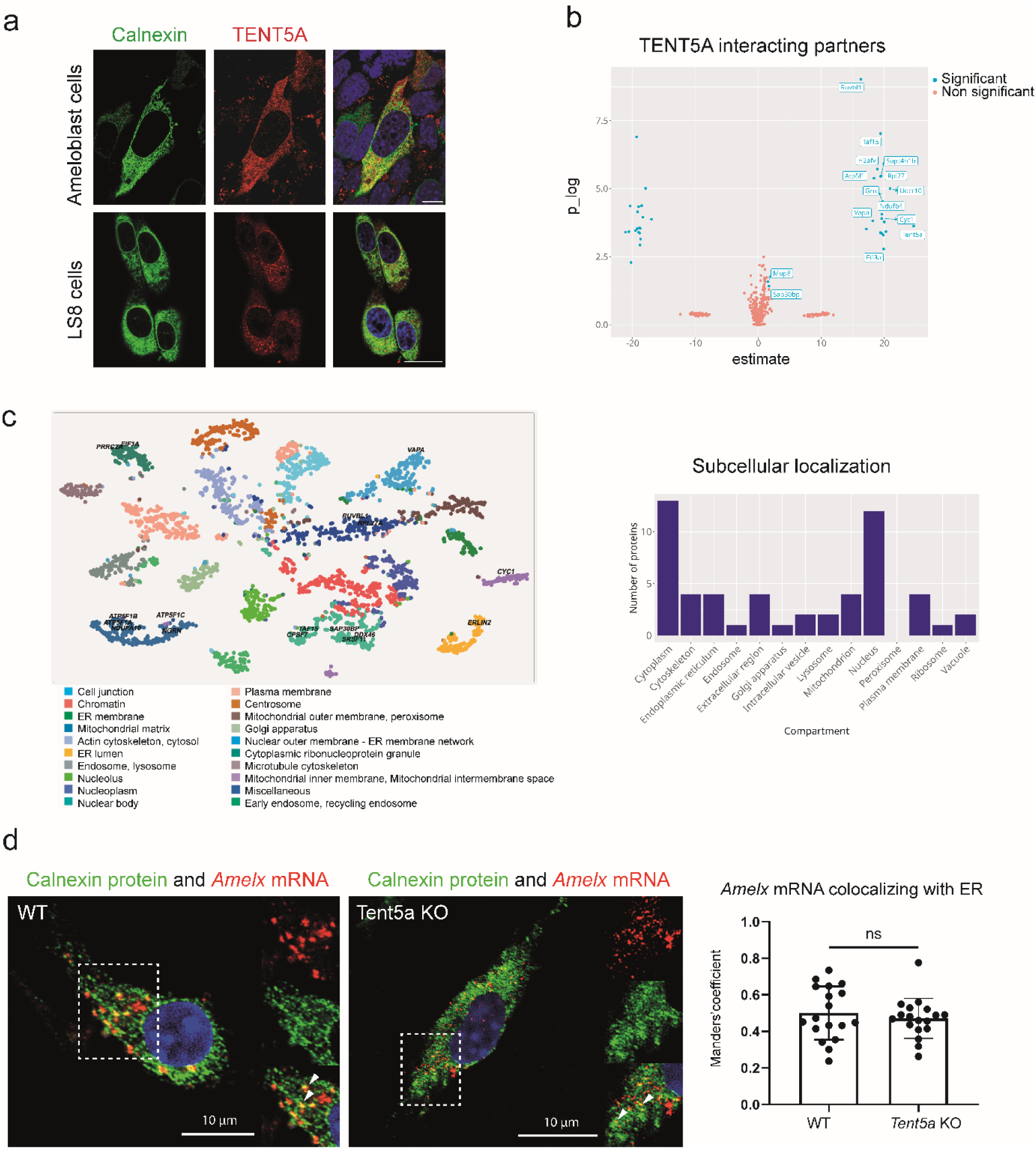
TENT5A subcellular localization and potential interacting partners identification. (**A**) Immunofluorescence staining of ameloblast immortalized cell line IMAL and LS8 cell line displays Calnexin ER protein (green) and Tent5a_Myc (red). Scale bar, 10μm. (**B**) Volcano plot mass spectrometry analysis of potential TENT5A interacting protein partners in ameloblast cells. P-value <0.03. (**C**) Subcellular localization of potential TENT5A interacting partners using Human cell map and subcellular compartment analysis from mass spectrometry data. P-value <0.03. (**D**) FISH assay showing *AmelX* and *Ambn* mRNA localization with Calnexin ER protein in WT and Tent5a KO ameloblasts primary cells. Scale bar, 10μm. Colocalization parameters were measured using Manders’ coefficient, statistical test Non-parametric Paired Wilcoxon test,p-value*<0.001.

TENT5A, along with other members of the TENT5 protein family, does not possess RNA binding motifs in its protein structure (*13*). A co-immunoprecipitation assay was performed to explore the potential formation of the TENT5A protein complex and its interaction with mRNA substrates. Mass spectrometry (MS) analysis identified various proteins, including ER proteins Granulin (GRN) and Erlin2. Additionally, associated with the role of TENT5A, EIF3A translation initiation protein was observed in the results as well as ribosome proteins such as RPL27. Notably, CPSF7, a cleavage and poly(A)denylation factor, was also identified, suggesting its significance for TENT5A function (Fig 5b). Then, TENT5A potential interacting partners were plotted by subcellular localization and cytoplasm, ER, and extracellular region were enriched (Fig 5c). Although many of the potential TENT5A binding partners identified in these experiments can be flagged as consistent with TENT5A biological function, it remains unclear whether any of these interactions assist in direct TENT5A function. Nonetheless, our findings confirmed TENT5A localization in the endoplasmic reticulum and its putative role in the translation initiation of a significant group of extracellular proteins.

To determine whether the localization of TENT5A substrate transcripts is directly influenced by TENT5A activity, we quantified the localization of *AmelX* mRNA using Calnexin ER protein staining. A portion of the mRNA was localized in the ER in both WT and Tent5a KO ameloblasts. There was no significant difference in the mRNA localization between genotypes, as evidenced by the Mander’s coefficient quantifying the colocalization of *AmelX* mRNA colocalizing with ER protein Calnexin (p value < 0.05) (Fig 5d). This suggests that the transport of transcripts to the ER membrane is largely independent of TENT5A activity.

In summary, our study demonstrates the crucial role of TENT5A in proper enamel formation. In ameloblasts, TENT5A is responsible for cytoplasmic polyadenylation of mRNAs encoding extracellular matrix proteins during the secretory stage. Among these proteins, AMELX, which is the primary component of the enamel organic matrix and essential for fully mineralized enamel formation, has been identified as a substrate of TENT5A adding another layer to the regulation of gene expression during amelogenesis. The downregulation of specific mRNAs, resulting from the removal of cytoplasmic polyadenylation in Tent5a KO mice, appears to be the underlying cause of the observed Amelogenesis imperfecta phenotype.

## Discussion

Using a Tent5a KO mouse model, we discovered that *Tent5a* deficiency leads to Amelogenesis imperfecta, highlighting its significance in enamel formation. μCT analysis revealed a reduction of the enamel layer and disruption of enamel ultrastructure. *Tent5a* is expressed in the cytoplasm of secretory stage ameloblasts in mouse incisors, consistent with previous studies showing its expression in developing tooth buds (*25*) and single-cell transcriptomics measuring the expression in distal secretory ameloblasts (*22*). Although its role in teeth has not been characterized until now, our results indicate that TENT5A primarily functions during amelogenesis by polyadenylating *AmelX* and other extracellular matrix transcripts, promoting their stability, translation and secretion.

Mutations in *Tent5a* can impair the expression of a larger set of genes involved in biomineralization process due to a shorter poly(A) tail, including *AmelX*, *Clu*, *Bgn* and *Col1a1*, resulting in reduced protein levels that contribute to AI development. The expression of *Ambn* and *Enam* was unaffected, suggesting that TENT5A specifically regulates the expression of selected secreted proteins in ameloblast cells. Notably, TENT5A polyadenylates specific AmelX isoforms, with Amelx-202 showing the most significant reduction in poly(A) tail length, making it a key TENT5A substrate. In contrast, Amelx-203, which has an additional exon, is not a TENT5A substrate. However, no specific recognition region responsible for substrate selection in *AmelX* mRNA has been identified. Furthermore, when exogenous Amelx mRNA was transfected into cells, there was a difference in mRNA stability between TENT5A substrates and non-substrate mRNA, but it was not dependent on the canonical poly(A) tail. This suggests that TENT5A does not need an existing poly(A) tail to identify the substrate to polyadenylate in the cytoplasm, indicating that substrate selection likely occurs during mRNA processing in nucleus.

While the downregulation of *AmelX* may be the main cause of the AI phenotype, the reduction of other extracellular matrix proteins also affects the physiology of the extracellular matrix, exacerbating the phenotype. For instance, BGN is thought to inhibit AMELX although the exact mechanism remains unclear (*26*). Then, another downregulated gene in Tent5a KO ameloblast is *Clu* and its expression is detected in mice molar tooth germs (*27*). Clusterin functions as a chaperone protein to stabilize extracellular misfolded proteins (*28*). Therefore, if CLU is downregulated in Tent5a KO mice, the tissue may struggle to regulate the folding of extracellular proteins like AMELX and AMBN. Lastly, while collagen is extensively studied in the context of bones, it is also expressed by ameloblast cells (*29*). Although its role in amelogenesis is not well understood, the overall dysregulation of extracellular transcripts affects the health of the extracellular organic matrix. Thus far, the ablation of *Clu, Bgn*, or *Col1a1* individually in mice has not been associated with a dental phenotype. However, the cumulative effect of dysregulated genes might contribute to the observed hypomineralization phenotype in Tent5a KO mice.

The consequences of gene dysregulation in the Tent5a KO mouse incisor were visualized by AMELX and AMBN immunostaining. The signals corresponding to these proteins were significantly reduced, and the synthesized proteins were unable to assemble in the extracellular matrix to form an organized organic matrix through AMELX-AMBN interactions. This disruption not only led to a decrease in the production of AMELX, but the synthesized AMELX and AMBN were secreted but not fully functional. *Tent5a* deletion resulted in changes in the AMELX-AMBN ratio within the organic enamel matrix, which is important for precise enamel formation (*30*). Both proteins undergo extensive intracellular and extracellular modifications, which are essential for proper self-assembly and enamel maturation (*31*).

Next, our study revealed the important role of TENT5A in the secretion and post-transcriptional regulation of extracellular matrix proteins. While we were unable to identify a consensus mRNA sequence for TENT5A substrates, we hypothesize that specificity may be determined by its protein binding partners and during RNA processing. Mass spectrometry analysis revealed several ER proteins and RNA-binding proteins that interact with TENT5A, suggesting its involvement in a larger regulatory complex. Additionally, our study confirmed the localization of TENT5A in the ER and investigated its function in exporting transcripts encoding extracellular matrix proteins to the ER. Notably, AmelX mRNA exhibited similar ER localization in both WT and Tent5a KO ameloblasts. Thus, TENT5A is crucial for maintaining the stability of AmelX mRNA and facilitating its translation in the ER, although it may not be essential for the export of transcripts to the ER.

Our findings align with previous studies demonstrating the role of TENT5A in bone formation by promoting the translation of collagen mRNA (*13*, *14*). Collectively, this study provides evidence that TENT5A is a key regulator of gene expression during amelogenesis, and its deficiency leads to amelogenesis imperfecta. It provides important insights into the molecular mechanisms underlying tooth development and highlights the critical role of TENT5A in post-transcriptionally regulating the expression of amelogenin and other secreted proteins for proper enamel tissue homeostasis.

## Materials and Methods

### Experimental Models

#### Mouse lines

*Tent5a* knock-out mouse line with a loss of function indel mutation in exon 2 (c.150_175del26bp – deletion of 26 bp starting at position 150 of cDNA – transcript ID ENSMUST00000034802.14) in C57BL/6N genetic background were established using the CRISPR/Cas9 method created for previous study (*14*).

#### Mice breeding conditions

Tent5a KO (C57BL/6N) mice were bred at the Czech Center of Phenogenomics and maintained in individually ventilated cages in a room with controlled temperature (22±2 °C) and humidity under a 12 h light/12 h dark cycle. Food (standard Altromin diet) and drink were provided *ad libitum*. The animals were closely followed by the animal caretakers and researchers, with regular inspection by a veterinarian, according to the standard animal welfare procedures of the local animal facility. All animal experiments were approved by the Animal Ethics Committee of the Czech Academy of Sciences (primary screen project number: 62/2016 and secondary screen project number: 45/2017) and were performed according to Czech guidelines for the Care and Use of Animals in Research and Teaching.

#### Cell lines

Immortalized ameloblast cell line was created by extracting the incisor teeth from 4-6 WT mice, soaked twice in 70% EtOH and transferred to HBSS with 1% StrepPen (15140122, Gibco). The cervical loop was extracted and incubated with 2 mg/ml collagenase type I (17101015, Gibco) and 3 mg/ml dispase (17105041, Gibco) in PBS, for digestion in an incubator at 37°C for 50 minutes under agitation. Then, the primary cell culture was incubated in a 6 well plate in DMEM/F12 media (12634010, Gibco) with 10% FBS (F7524, Sigma-Aldrich) and 1% StrepPen. After 24h, the dish was washed with PBS and attached cells were grown for 10 days. Ameloblast looking like colony was immortalized with the SV40 Cell Immortalization System using the following standard protocol (*32*) with plasmids pLenti CMV/TO SV40 small + Large T (w612-1) (22298, Addgene), pCMV-VSV-G (8454, Addgene) and pCMV-dR8.2 dvpr (8455, Addgene). Immortalized cells were separated using flow cytometry to get monoclonal cell lines that were quantified by RT-qPCR for enamel proteins.

Ameloblast like LS8 mouse cell line (*24*) were gifted from INSERM (French National Institute of Health and Medical Research), Strasbourg, France.

#### Plasmids

Salivary gland RNA was converted into cDNA with M-MLV Reverse Transcriptase (A3500, Promega) using random primers to use it as a template for *Tent5a* CDS synthesis with 2x Phusion master mix (F531L, Thermo Scientific). MluI and AsiSI restriction enzyme sites were added with Forward: 5’-TAGCGATCGCCATGGCCGAGGGCGAAGGGTA-3’ and Reverse: 5’-TTACGCGT GTTGCAGGGTAACCAAGTAGAGTATG – 3’ primers and used for ligation with T4 ligase (M1801, Promega) to pCMV6-Entry (PS100001, Origene). Tent5a_Myc plasmid was sequenced using T7 promoter primer to confirm *Tent5a* gene insertion.

Luc2 sequence was cloned from pGL4.10[luc2] to pGEM-T Easy plasmid (A1360, Promega) to create luc2_pGEM. ENSMUST00000112118.8 (Amelx-202) cDNA from ameloblast cells was used to clone Amelx-202 into a pGEM-T Easy plasmid (A1360, Promega). ENSMUST00000112119.8 (Amelx-203) was generated by adding an exon to Amelx-202 pGEM by In-Fusion cloning (639648, Takara Bio) using primers:

Forward: TCAGTATTGATAGCCTGAGAATGTGACTTCTCATAGCTTAAGTTGATATAACCA

Reverse: GGCTATCAATACTGACAGGACTGCATTAGTGCTTACCCCTTTGAAGTGGT

#### RNA in situ hybridization

*Tent5a* RNA probes were generated from PCR fragment amplified with primers Forward: 5’-GACCGCTGCTCGGACTATTG- 3’ and Reverse: 5’- GTCTTCCAAGCCCACGAAGT- 3’ and cloned in pGEM-T Easy (A1360, Promega) following manufactures instructions. The digoxygenin-labeled antisense and sense RNA probes were synthesised with the RNA DIG labelling kit (11277065910, Roche) by in vitro transcription. Sense probes were used for negative control samples.

Mice were sacrificed and heart perfusion with 4% paraformaldehyde (PFA) was used to fixate. The mandibles were extracted and fixed in 4% PFA for 24 h at 4 °C. Then, they were placed in Morse’s solution for decalcification (22.5% Formic acid and 10% sodium citrate) for 3 days, dehydrated, and mounted in paraffin. 7 μm thick sections were sectioned in an RNAse-free environment. In situ hybridization was performed according to a standard protocol (*33*). RNA probes were detected by 1μl/5ml anti-digoxigenin antibody (11093274910, Roche) and BM-purple (11442074001, Roche) was used to detect RNA according to the manufacturer’s instructions. The images were obtained with Zeiss Axioscan Z2 (Carl Zeiss).

#### Micro-computed tomography scanning and analysis of teeth (full body and teeth HR Phase contrast)

First, the skull scan was performed on a SkyScan 1176 instrument (Bruker) at a resolution of 9 µm per voxel (0,5 mm Al filter; voltage, 50 kV; current, 500 µA; exposure, 900 ms; rotation, 0,3°; spiral scan, 2x averaging) in a wet atmosphere. Reconstruction was performed in an NRecon 1.7.1.0 (Bruker) with the following parameters: smoothing = 3, ring artifact correction = 3, beam hardening correction = 0%, and defect pixel masking threshold = 0%. The intensities were set from 0,004 AU to 0,123 AU.

Then, we proceed with high-resolution MicroCT (HR-μCT) scanning in SkyScan 1272 High-Resolution X-Ray Microtomograph (Bruker). Incisors were extracted and embedded in 2.5% low-temperature gelling agarose (Lonza). The sample was left in a refrigerator (4 °C) for sample stabilization for 1 day. Incisor teeth were scan at a resolution of 1.5 μm per voxel (Al filter, 1 mm; voltage, 80 kV; current, 125 mA; exposure, 3100 ms). Proximal and distal borders were selected for scanning the teeth: 360° scan, rotation step, 0,21°, 2x averaging. Single-distance phase retrieval was performed at Delta/beta parameter at 370 for the phase contrast. The reconstruction was done in NRecon 2.0.0.5 at smoothing=5, ring artifact reduction= 14, beam hardening correction= 28%, and defect pixel masking threshold=10%. The intensities were set from 0,00 AU to 0,12 AU.

#### Incisor enamel visualization in SEM

Lower incisors of mice were extracted, and the distal tip was removed using scissors. Inside the melted epon resin (Embed 812 kit) incisors were embedded in a longitudinal orientation and put in a 60 °C incubator for resin polymerization for 3 days. The tip of the incisor was ground sagittally and etched with 3 % HCl for 5s. Finally, colloidal silver was used to decrease the charging. The samples were dried overnight, and finally sputter-coated with 10 nm platinum.

The electron microscope was used Nova NanoSEM 450 scanning electron microscope (FEI) at 5 kV or Dual beam system FEI Helios NanoLab 660 G3 UC at 1 kV and 0,1 nA using an Everhart-Thornley secondary electron detector secondary electron mode (SE). The SEM acquisition was performed in the Imaging Methods Core Facility at BIOCEV.

#### Histology and Immunohistochemistry

Mice were euthanized by cervical dislocation and heart perfused with 4% PFA. Mandibles were extracted and fixed in 4% paraformaldehyde at 4°C for 48h followed by decalcification in Osteosoft (1017281000, Sigma-Aldrich) for 14 days. Samples were embedded in paraffin using a Leica EG 1150H paraffin embedding station (Leica Microsystems) and cut into 6 μm thick sections, deparaffinized, and rehydrated.

Hematoxylin eosin (H&E) staining was performed in Leica automated stainer (Leica ST5020 + Leica ST5030). The images were captured using a Zeiss Axio imager Z2 microscope (Carl Zeiss).

For immunohistochemistry, antigen retrieval was performed for 10’ in the pressure cooker with HIER EDTA Buffer pH 8.0 (ZUC040, Zytomed). The samples were blocked by 2.5% normal goat serum (S-1012, Vector Laboratories) for 1h at room temperature and incubated overnight with primary antibodies against Ambn (diluted 1:50, AF3026, R&D Systems) or AmelX (diluted 1:50, ABT260 Sigma-Aldrich). After washing with PBS, slides were incubated with the secondary antibody Alexa fluor 488 and 594 (diluted 1:1000, Thermo Fisher Scientific) for 1h at room temperature. DAPI (1:1000) was used as counterstain. Images were captured using Spinning disk Nikon Eclipse Ti2 (Yokogawa CSU-W1) confocal microscopy.

#### Immunocytochemistry

Ameloblast immortalized cells and LS8 cells were transfected with Tent5a_Myc using Lipofectamine 2000 (Invitrogen). After 48h, I washed the cells with PBS and fixed with 4% PFA for 20’ at room temperature. Then, I performed immunocytochemistry with primary antibody Calnexin (1:100, ab22595, Abcam) and Myc (1:100, ab32, Abcam) overnight at 4°C. After washing 3 times with PBS, slides were incubated with the secondary antibody Alexa fluor 488 and 594 (diluted 1:500, Thermo Fisher Scientific) for 1h at room temperature. Next, the cells were counterstain with DAPI 1:1000 and mount in fluorescent mounting medium (S302380, Dako). Images were obtained in a Spinning disk Nikon Eclipse Ti2 (Yokogawa CSU-W1) confocal microscopy.

#### mRNA-FISH and Immunofluorescence

Stellaris single molecule FISH probes were purchased from BioSearch Technologies, Ambn Quasar 570 and Amelx Quasar 670. LS8 cells were washed with PBS and fixed for 10’ in 4% PFA, permeabilized in 70% ethanol overnight at 4 °C. Cells were washed twice with PBS and blocked for 30’ with 5% nuclease free BSA (126615, Sigma-Aldrich). Then, cells were stained with primary antibody for 2h (1:100 Myc, Ab32, Abcam or 1:100 Calnexin, ab22595, Abcam) at room temperature diluted in 1% BSA in PBS. Cells were rinsed with wash buffer A (SMF-WA1-60, BioSearch technologies) and hybridized with RNA probe diluted in hybridization buffer (1:100, SMF-HB1-10, BioSearch technologies) overnight at 37°C in a humified chamber. Then, the coverslips were washed with wash buffer A and corresponding secondary antibody Alexa Fluor (1:500, Thermo Fisher Scientific) for 30’ at 37°C. After, samples were washed with wash buffer A and DAPI (5ng/ml) for 30’ at 37°C. Coverslips were washed with wash buffer B (SMF-WB1-20, BioSearch technologies) for 5’ and mounted in fluorescent mounting (S302380, Dako). Images were taken in a Spinning disk Nikon Eclipse Ti2 (Yokogawa CSU-W1) confocal microscopy with 100% laser power for FISH signal. Colocalization was measured in ImageJ BIOP Jacop plugin (n=18) and statistical analysis of Mander’s coefficient was tested with Non-parametric Paired Wilcoxon test.

#### RNA isolation

Mandibles were isolated and kept in RNA later (Thermofisher Scientific) for 24-48h. The tissues were mechanically ground with IKA T10 basic Ultra-Turrax (IKA) and RNA was isolated using the RNeasy mini kit (74106, Qiagen) with DNase kit (79254, Qiagen) following manufacturer’s instructions.

#### RT-qPCR

RNA was used as a template for reverse transcription into cDNA with M-MLV Reverse Transcriptase (M1701 and M5313, Promega) using random primers (C1181, Promega). Quantitative PCR (qPCR) reactions were performed using the Light Cycler® 480 SYBR® Green I Master (4887352001, Roche) in Light Cycler® 480 Instrument II (Roche). Primers were designed and ordered from Sigma **(*Ambn*** F: 5’-ccaggttgttgaggaaatgc-3‘, R: 5‘-cacagtgaatgtcagcatctaag-3’; ***AmelX*** F: 5’-gcatacactcaaagaaccatcaag-3’, R: 5‘-cacctcatagcttaagttgatataacc--3’; ***Mmp20*** F: 5’-gctaactaccgcctcttcc-3’, R: 5’-ccatctgtattgccttgtcca-3’; ***Clu*** F: 5’-gtccagggagtgaagcacat-3’, R: 5‘-tccctagtgtcctccagagc-3’; ***Enam*** F: 5’-tgcagaaatccgacttctcct-3’, R: 5’-catctggaatggcatggca-3’; ***Col1a1*** F: 5’-cctcagggtattgctggacaac-3’, R: 5’-cagaaggaccttgtttgccagg-3’; ***Bgn*** F: 5’- ggctactcaccttgctgctg-3’, R: 5’- gagcagcccatcatccaagg -3’; ***Tent5a*** F: 5’- ggggaagaggagtttcagac -3’, R: 5’-gcacataagcctccttgaga -3’). The expression levels of the genes of interest were normalized to the levels of ***Rpl19*** (F: 5’-aagcctgtgactgtccattc-3’, R: 5’-gatcctcatccttctcatccag-3’). All experiments were performed independently in triplicates on 4-5 WT and Tent5a KO specimens. Graphs were generated using GraphPad Prims version 8.0.0 and student t-test *p-value p<0.05, **p-value p<0.01, ***p-value p<0.001, ****p-value p<0.0001 was used for statistical significance.

#### Illumina mRNA sequencing

RNA from 3 wild-type and 3 Tent5a KO cervical loop was analyzed for quality and 1μg of RNA was sent for library preparation with poly(A) selection (Lexogen kit) and library preparation (SENSE Total RNA-Seq Library Prep Kit). Sequencing that included pool preparation and quality control (Agilent High Sensitivity DNA), NextSeq PhiX Control Kit and mRNA sequencing was performed with NextSeq® 500/550 High Output Kit v2 (75 cycles) (Illumina, USA). The results were analyzed for QC using MultiQC tool (https://multiqc.info/) and then there were analyzed by standard way using DESeq2 library (https://bioconductor.org/packages/release/bioc/html/DESeq2.html). Data was visualized in R (www.R-project.org).

#### *Nanopore sequencing-* Nanopore direct RNA sequencing (DRS)

Direct RNA sequencing was performed as described by (*34*). Briefly, RNA libraries were prepared from 500 ng murine cap-enriched mRNA mixed with 50-200 ng oligo-(dT)_25_-enriched mRNA from *Saccharomyces cerevisiae* yeast with a Direct RNA Sequencing Kit (catalog no. SQK-RNA002, Oxford Nanopore Technologies) according to the manufacturer’s instructions. Sequencing was performed using R9.4 flow cells on a MinION device (ONT). Raw data were basecalled using Guppy (ONT). Raw sequencing data (fast5 files) were deposited at the European Nucleotide Archive (ENA, accession numbers to be provided).

#### Poly(A) lengths determination and statistical analysis

Mouse-originating DRS reads were separated from the other samples by mapping to the Gencode VM26 transcriptome using Minimap2 2.17 with options -k 14 -ax map-ont – secondary=no and processed with samtools 1.9 to filter out supplementary alignments and reads mapping to reverse strand (samtools view -b -F 2320). The poly(A) tail lengths for each read were estimated using the Nanopolish 0.13.2 polya function^25^. In subsequent analyses, only length estimates with the QC tag that was reported by Nanopolish as PASS were considered. Statistical analysis was performed using functions provided in the NanoTail R package (https://github.com/smaegol/nanotail, manuscript in preparation). In detail, the Generalized Linear Model approach, with log2(polya length) as a response variable, was employed, and transcripts that had a low number of supporting reads in each condition (<20) were filtered out. To correct for the batch effect, a replicate identifier was used as one of the predictors, in addition to the condition (Tent5a KO/WT) identifier. P values (for the condition effect) were estimated using the Tukey HSD post hoc test and adjusted for multiple comparisons using the Benjamini–Hochberg method. Transcripts were considered as having a significant change in poly(A) tail length, if the adjusted P value was < 0,05, and there were at least 20 supporting reads for each condition.

#### PAT assay

PAT assay was performed using the Poly (A) Tail-Length Assay Kit (764551kt, Thermo scientific) following manufacturer’s instructions. The primers used were the following: *AmelX* F: 5‘-CAGTCACCTCTGCATCCCAT-3’ and R: 5’-TCCATGTTAAGCGGATGCCT-3’, *Clu* F: 5’-AAGGCGCTACAGGAATACCG-3’, and R: 5’-TTCTTCCCGAGAGCAGCAAG-3’ and *Ambn* F: 5’-CAGCCACTGCTACCTGGAAA-3’ and R: 5’-GGAAGCAAGAAGGGACCTACA-3’. 1μg of wild-type RNA and Tent5a KO cervical loop was used for the starting reaction. The PCR product was analyzed on a 2.5% agarose gel.

#### mRNA transfection

Amelx-202_pGEM, Amelx-203_pGEM and luc2_pGEM were used for in vitro transcription using mMESSAGE mMACHINE T7 Transcription Kit (AM1344, Invitrogen) following manufacturers’ instructions. A poly(A) tail was added to the transcribed mRNA using Poly(A) Tailing Kit (AM1350, Invitrogen) and mRNA tailing was validated in an agarose gel.

Ameloblast immortalized cells were plated and 25ng of mRNA was transfected using Lipofectamine™ MessengerMAX™ Transfection Reagent (LMRNA008, Invitrogen). RNA was extracted at different timepoints (3h, 6h and 20h) to measure mRNA levels using RT-PCR. Primers for RT-qPCR were the following: *Amelx 202* F: 5‘-AGGGGTAAGCACCTCATAGCTTA -3’ and R: 5’-GGGGTAAGCACCTCATAGCTTA -3’, *Amelx 203* F: 5‘-TGGATTTTGTTTGCCTGCCT-3’ and R: 5’-TGATAGCCTGAGAATGTGACTTC-3’, and *luciferase* F: 5‘- GCTACAAACGCTCTCATCGACAAG -3’ and R: 5’- GTATTTGATCAGGCTCTTCAGCCG -3’.

#### Western blot

For teeth samples western blot analysis, incisor tips were cut out and the remaining incisors from two mice were combined for protein extraction in 200μl 1% NP40 lysis buffer complemented with protease inhibitors. Teeth were homogenized with IKA T10 basic Ultra-Turrax (IKA) and incubated on ice for 30 min inverting the tube. Then, samples were spun at 12000rpm at 4°C for 20 min and the supernatant was used for protein concentration analysis (Pierce BCA Protein Assay kit, Thermo Fisher Scientific). 5μg of protein were resolved on a 4–20% Mini-PROTEAN TGX Precast Protein Gel (Biorad) and were electro-transferred onto nitrocellulose membranes (Biorad). Blocking was performed in 5% milk in TBS-T for 1h, followed by antibody incubation overnight at 4°C, anti-AMBN (1:1000, AF3026, R&D), Anti-AMELX 1:5000 (ABT260, Sigma), Anti-Clu 1:3000 (AF2747, R&D), Anti Vinculin as a housekeeping gene (1:5000, ab129002, Abcam) diluted in TBS with 3% BSA and 0.1% azide. After washing with TBS-T, secondary antibody incubation (1:10000) was done with Mouse anti-rabbit (sc-2030, Santa Cruz) and Mouse anti-goat IgG-HRP (ab97110, Abcam) for 1h at room temperature. Gels were visualized with SuperSignal™ West Pico PLUS Chemiluminescent Substrate Kit (Thermo Fisher Scientific).

For Co-IP samples immunoblotting, Tent5a was detected in the membrane with Anti c-Myc (1:3000, C3956, Sigma-Aldrich), Histone-H3 (1:5000, ab1791, Abcam) as a nuclear fraction reference and Anti Vinculin as a cytoplasmatic reference.

#### Co-Immunoprecipitation

IMAL Ameloblast cells were transfected with Tent5a_Myc plasmid and 48h later, cells were lysed for nuclear and cytoplasmatic fractionation following a standard protocol (*35*). Shortly, PBS was added, and cells were scraped and centrifuged at 980 rpm for 4’. Then, cell lysis buffer [10mM HEPES; pH7.5, 10mM KCl, 0.1mM EDTA, 1mM dithiothreitol (DTT), 0.5% Nonidet-40 and 0.5 mM PMSF] complemented with protease inhibitors (PhosSTOP 4906845001 and cOmplete 11697498001, Roche) was added to the pellet. Cells were resuspended and incubated for 30 minutes in ice with intermittent mixing. Cells were collected by centrifuging at 12000g at 4°C for 15’. The supernatant was stored as the cytoplasmic fraction. Then, the pellet was incubated with 100μl of nuclear lysis buffer [20mM HEPES; pH7.5, 400mM NaCl, 1mM EDTA, 1mM DTT and 1mM PMSF] with protease inhibitors and during this time, sonicated for 2 seconds. Centrifugation was used (12000g 15’ at 4°C) to collect the nuclear fraction.

Cells without the Ten5a_Myc plasmid were used as a negative control. Pierce c-Myc Tag IP/Co-IP Kit (23620, Thermo scientific) was used for co-immunoprecipitation and was performed according to the manufacturer’s protocol. Proteins were analyzed in mass spectrometry for protein detection.

#### Protein Digestion- Mass spectrometry

LC-MS analyses were performed in Laboratory of Mass Spectrometry at Biocev research center, Faculty of Science, Charles University. Columns were washed by 100mM TEAB containing 2% SDC. Cysteins were reduced with 10mM final concentration of TCEP and blocked with 40mM final concentration of chloroacetamide (60°C for 30 min). Samples were cleaved on beads with 1µg of trypsin at 37°C overnight. After digestion samples were centrifuged and supernatants were collected and acidified with TFA to 1% final concentration. SDC was removed by extraction to ethylacetate (Masuda et al, 2008). Peptides were desalted using in-house made stage tips packed with C18 disks (Empore) (*36*).

#### nLC-MS 2 Analysis- Mass spectrometry

Nano Reversed-phase column (EASY-Spray column, 50 cm x 75 µm ID, PepMap C18, 2 µm particles, 100 Å pore size) was used for LC/MS analysis. Mobile phase buffer A was composed of water and 0.1% formic acid. Mobile phase B was composed of acetonitrile and 0.1% formic acid. Samples were loaded onto the trap column (Acclaim PepMap300, C18, 5 µm, 300 Å Wide Pore, 300 µm x 5 mm, 5 Cartridges) for 4’ at 15μl/min. Loading buffer was composed of water, 2% acetonitrile and 0.1% trifluoroacetic acid. Peptides were eluted with Mobile phase B gradient from 4% to 35% B in 60 min. Eluting peptide cations were converted to gas-phase ions by electrospray ionization and analyzed on a Thermo Orbitrap Fusion (Q-OT-qIT, Thermo). Survey scans of peptide precursors from 350 to 1400 m/z were performed at 120K resolution (at 200 m/z) with a 5 × 10^5^ ion count target. Tandem MS was performed by isolation at 1,5 Th with the quadrupole, HCD fragmentation with normalized collision energy of 30, and rapid scan MS analysis in the ion trap. The MS/MS ion count target was set to 10^4^ and the max injection time was 35ms. Only those precursors with charge state 2–6 were sampled for MS/MS. The dynamic exclusion duration was set to 45 s with a 10ppm tolerance around the selected precursor and its isotopes. Monoisotopic precursor selection was turned on. The instrument was run in top speed mode with 2 s cycles (*37*).

#### Data analysis- Mass spectrometry

All data were analyzed and quantified with the MaxQuant software (version 2.0.2.0) (*38*). The false discovery rate (FDR) was set to 1% for both proteins and peptides and we specified a minimum peptide length of seven amino acids. The Andromeda search engine was used for the MS/MS spectra search against the *Mus musculus* database (downloaded from uniprot.org, containing 17 059 entries). Enzyme specificity was set as C-terminal to Arg and Lys, also allowing cleavage at proline bonds and a maximum of two missed cleavages. Carbmaidomethylation of cysteine was selected as fixed modification and N-terminal protein acetylation and methionine oxidation as variable modifications. The “match between runs” feature of MaxQuant was used to transfer identifications to other LC-MS/MS runs based on their masses and retention time (maximum deviation 0.7 min) and this was also used in quantification experiments. Quantifications were performed with the label-free algorithms described recently. Data analysis was performed using Perseus 1.6.15.0 software.

#### Statistical analysis

The statistical tests utilized are specified in each method. Normality was evaluated using the Shapiro-Wilk test for one-way ANOVA, two-way ANOVA, and unpaired two-tailed t-tests. Statistical analysis and graph plotting were conducted using Prism v10.2.3 (GraphPad).

## Acknowledgements

The authors acknowledge Imaging Methods Core Facility at BIOCEV for their support & assistance in this work. The authors also acknowledge the OMICS Proteomics BIOCEV core facility for its excellent technical service.

## Funding

The author(s) declare financial support was received for the research, authorship, and/or publication of this article.

Czech Science Foundation CSF Grant 19-19025Y

Czech Academy of Sciences RVO 68378050 and by grants

Ministry of Education, Youth and Sports of the Czech Republic LM2018126, LM2023050 and LM2023036

European Structural and Investment Funds CZ.02.1.01/0.0/0.0/18_046/0015861

European Research Council ERC AdG 101097317

European Funds for Smart Economy 2021–2027 (FENG)

## Author contributions

GAN and JP designed and conducted the experiments. OG, SM, PK, and AD carried out direct RNA sequencing. FS and GA performed micro-CT scanning and analysis, while VN and CMT analyzed the transcriptomics and mass spectrometry data. PT and KH conducted protein mass spectrometry. AAV performed the mRNA transfection experiment. IK and AB carried out the SEM experiment. JP, RS, and AD supervised the experiments. GA wrote the manuscript, with contributions from JP, FS, OG, AD, and RS.

## Competing interests

Authors declare no conflict of interests.

## Notes

### Competing Interest Statement

The authors have declared no competing interest.

## References

1. L. Weill, E. Belloc, F. A. Bava, R. Méndez, Translational control by changes in poly(A) tail length: Recycling mRNAs. Nat Struct Mol Biol 19, 577–585 (2012).

2. S. Yu, V. N. Kim, A tale of non-canonical tails: gene regulation by post-transcriptional RNA tailing. Nat Rev Mol Cell Biol 21, 542–556 (2020).

3. C. Y. A. Chen, A. Bin Shyu, Mechanisms of deadenylation-dependent decay. Wiley Interdiscip Rev RNA 2, 167–183 (2011).

4. C. J. Norbury, Cytoplasmic RNA: A case of the tail wagging the dog. Nat Rev Mol Cell Biol 14, 643–653 (2013).

5. T. Nakanishi, H. Kubota, N. Ishibashi, S. Kumagai, H. Watanabe, M. Yamashita, S. I. Kashiwabara, K. Miyado, T. Baba, Possible role of mouse poly(A) polymerase mGLD-2 during oocyte maturation. Dev Biol 289, 115–126 (2006).

6. L. Yan, M. Yang, H. Guo, L. Yang, J. Wu, R. Li, P. Liu, Y. Lian, X. Zheng, J. Yan, J. Huang, M. Li, X. Wu, L. Wen, K. Lao, R. Li, J. Qiao, F. Tang, Single-cell RNA-Seq profiling of human preimplantation embryos and embryonic stem cells. Nat Struct Mol Biol 20, 1131–1139 (2013).

7. K. W. Kim, T. L. Wilson, J. Kimble, GLD-2/RNP-8 cytoplasmic poly(A) polymerase is a broad-spectrum regulator of the oogenesis program. Proc Natl Acad Sci U S A 107, 17445–17450 (2010).

8. Z. Warkocki, V. Liudkovska, O. Gewartowska, S. Mroczek, A. Dziembowski, Terminal nucleotidyl transferases (TENTs) in mammalian RNA metabolism. Philosophical Transactions of the Royal Society B: Biological Sciences 373 (2018).

9. L. Rouhana, L. Wang, N. Buter, E. K. Jae, C. A. Schiltz, T. Gonzalez, A. E. Kelley, C. F. Landry, M. Wickens, Vertebrate GLD2 poly(A) polymerases in the germline and the brain. Rna 11, 1117–1130 (2005).

10. S. Mroczek, J. Chlebowska, T. M. Kuliński, O. Gewartowska, J. Gruchota, D. Cysewski, V. Liudkovska, E. Borsuk, D. Nowis, A. Dziembowski, The non-canonical poly(A) polymerase FAM46C acts as an onco-suppressor in multiple myeloma. Nat Commun 8 (2017).

11. M. Doyard, S. Bacrot, C. Huber, M. Di Rocco, A. Goldenberg, M. S. Aglan, P. Brunelle, S. Temtamy, C. Michot, G. A. Otaify, C. Haudry, M. Castanet, J. Leroux, J. P. Bonnefont, A. Munnich, G. Baujat, P. Lapunzina, S. Monnot, V. L. Ruiz-Perez, V. Cormier-Daire, FAM46A mutations are responsible for autosomal recessive osteogenesis imperfecta. J Med Genet 55, 278–284 (2018).

12. I. Barragán, S. Borrego, M. M. Abd El-Aziz, M. F. El-Ashry, L. Abu-Safieh, S. S. Bhattacharya, G. Antiñolo, Genetic analysis of FAM46A in Spanish families with autosomal recessive retinitis pigmentosa: Characterisation of novel VNTRs. Ann Hum Genet 72, 26–34 (2008).

13. K. Kuchta, A. Muszewska, L. Knizewski, K. Steczkiewicz, L. S. Wyrwicz, K. Pawlowski, L. Rychlewski, K. Ginalski, FAM46 proteins are novel eukaryotic non-canonical poly(A) polymerases. Nucleic Acids Res 44, 3534–3548 (2016).

14. O. Gewartowska, G. Aranaz-Novaliches, P. S. Krawczyk, S. Mroczek, M. Kusio-Kobiałka, B. Tarkowski, F. Spoutil, O. Benada, O. Kofroňová, P. Szwedziak, D. Cysewski, J. Gruchota, M. Szpila, A. Chlebowski, R. Sedlacek, J. Prochazka, A. Dziembowski, Cytoplasmic polyadenylation by TENT5A is required for proper bone formation. Cell Rep 35 (2021).

15. J. Y. Sire, T. Davit-Béal, S. Delgado, X. Gu, The origin and evolution of enamel mineralization genes. [Preprint] (2007). 10.1159/000102679.

16. T. Martin, “Incisor enamel microstructure and systematics in rodents” in Tooth Enamel Microstructure, W. V. Koenigswald, Sander P. M., Eds. (Balkema, Rotterdam, 1997), pp. 163–175.

17. T. Wald, A. Osickova, M. Sulc, O. Benada, A. Semeradtova, L. Rezabkova, V. Veverka, L. Bednarova, J. Maly, P. Macek, P. Sebo, I. Slaby, J. Vondrasek, R. Osicka, Intrinsically disordered enamel matrix protein ameloblastin forms ribbon-like supramolecular structures via an N-terminal segment encoded by exon 5. Journal of Biological Chemistry 288, 22333–22345 (2013).

18. K. Kawasaki, K. M. Weiss, Mineralized tissue and vertebrate evolution: The secretory calcium-binding phosphoprotein gene cluster. Proceedings of the National Academy of Sciences 100, 4060–4065 (2003).

19. P. A. Fang, J. F. Conway, H. C. Margolis, J. P. Simmer, E. Beniash, Hierarchical self-assembly of amelogenin and the regulation of biomineralization at the nanoscale. Proceedings of the National Academy of Sciences 108, 14097–14102 (2011).

20. C. E. L. Smith, J. A. Poulter, A. Antanaviciute, J. Kirkham, S. J. Brookes, C. F. Inglehearn, A. J. Mighell, Amelogenesis imperfecta; genes, proteins, and pathways. Front Physiol 8 (2017).

21. P. J. Aldred, M. J., Savarirayan, R., & Crawford, Amelogenesis imperfecta: a classification and catalogue for the 21st century. Oral Dis 9, 19–23 (2003).

22. A. Sharir, P. Marangoni, R. Zilionis, M. Wan, T. Wald, J. K. Hu, K. Kawaguchi, D. Castillo-Azofeifa, L. Epstein, K. Harrington, P. Pagella, T. Mitsiadis, C. W. Siebel, A. M. Klein, O. D. Klein, A large pool of actively cycling progenitors orchestrates self-renewal and injury repair of an ectodermal appendage. Nat Cell Biol 21, 1102–1112 (2019).

23. R. A. Bapat, J. Su, J. Moradian-Oldak, Co-Immunoprecipitation Reveals Interactions Between Amelogenin and Ameloblastin via Their Self-Assembly Domains. Front Physiol 11 (2020).

24. L. S. Chen, R. I. Couwenhoven, D. Hsu, W. Luo, M. L. Snead, Maintenance of amelogenin gene expression by transformed epithelial cells of mouse enamel organ. Arch Oral Biol 37, 771–778 (1992).

25. G. E. Etokebe, A. M. Küchler, G. Haraldsen, M. Landin, H. Osmundsen, Z. Dembic, Family-with-sequence-similarity-46, member A (Fam46a) gene is expressed in developing tooth buds. Arch Oral Biol 54, 1002–1007 (2009).

26. M. Goldberg, D. Septier, O. Rapoport, M. Young, L. Ameye, Biglycan is a Repressor of Amelogenin Expression and Enamel Formation: An Emerging Hypothesis. J Dent Res 81, 520–524 (2002).

27. Q. E. S. Khan, A. Sehic, C. Khuu, S. Risnes, H. Osmundsen, Expression of clu and Tgfb1 during murine tooth development: Effects of in-vivo transfection with anti-miR-214. Eur J Oral Sci 121, 303–312 (2013).

28. S. Satapathy, M. R. Wilson, The Dual Roles of Clusterin in Extracellular and Intracellular Proteostasis. Elsevier Ltd [Preprint] (2021). 10.1016/j.tibs.2021.01.005.

29. N. Assaraf-Weill, B. Gasse, J. Silvent, C. Bardet, J. Y. Sire, T. Davit-Béal, Ameloblasts express type I collagen during amelogenesis. J Dent Res 93, 502–507 (2014).

30. M. L. Paine, H. J. Wang, W. Luo, P. H. Krebsbach, M. L. Snead, A transgenic animal model resembling amelogenesis imperfecta related to ameloblastin overexpression. Journal of Biological Chemistry 278, 19447–19452 (2003).

31. C. Robinson, J. Kirkham, S. J. Brookes, W. A. Bonass, R. C. Shore, The chemistry of enamel development. Int J Dev Biol 39, 145–52 (1995).

32. M. J. Tevethia, Immortalization of primary mouse embryo fibroblasts with SV40 virions, viral DNA, and a subgenomic DNA fragment in a quantitative assay. Virology 137, 414–421 (1984).

33. D. G. Wilkinson, M. A. Nieto, [22] Detection of Messenger RNA by in Situ Hybridization to Tissue Sections and Whole Mounts. Methods Enzymol 225, 361–373 (1993).

34. A. Bilska, M. Kusio-Kobiałka, P. S. Krawczyk, O. Gewartowska, B. Tarkowski, K. Kobyłecki, D. Nowis, J. Golab, J. Gruchota, E. Borsuk, A. Dziembowski, S. Mroczek, Immunoglobulin expression and the humoral immune response is regulated by the non-canonical poly(A) polymerase TENT5C. Nat Commun 11, 1–17 (2020).

35. E. Schreiber, K. Harshman, I. Kemler, U. Malipiero, W. Schaffner, A. Fontana, Astrocytes and glioblastoma cells express novel octamer-DNA binding proteins distinct from the ubiquitous Oct-1 and B cell type Oct-2 proteins. Nucleic Acids Res 18, 5495–5503 (1990).

36. J. Rappsilber, M. Mann, Y. Ishihama, Protocol for micro-purification, enrichment, pre-fractionation and storage of peptides for proteomics using StageTips. Nat Protoc 2, 1896–1906 (2007).

37. A. S. Hebert, A. L. Richards, D. J. Bailey, A. Ulbrich, E. E. Coughlin, M. S. Westphall, J. J. Coon, The one hour yeast proteome. Molecular and Cellular Proteomics 13, 339–347 (2014).

38. J. Cox, M. Y. Hein, C. A. Luber, I. Paron, N. Nagaraj, M. Mann, Accurate proteome-wide label-free quantification by delayed normalization and maximal peptide ratio extraction, termed MaxLFQ. Molecular and Cellular Proteomics 13, 2513–2526 (2014).

